# Mechanical energetic contributions of the rectus femoris during perturbed walking

**DOI:** 10.1101/2025.08.20.671067

**Authors:** Pawel R. Golyski, Gregory S. Sawicki

## Abstract

Animals maintain locomotor stability following external perturbations by coordinating muscular responses to produce desired mechanical behavior at different levels of description (*e*.*g*., muscle-tendon units, joints, legs). To investigate the role of proximal musculature in responding to perturbations during human walking, here we extend a previous analysis relating joint and leg levels down to the level of the rectus femoris. Using *in-vivo* B-mode ultrasound processed with a custom automated fascicle tracking application and EMG measurements to drive Hill-type models of muscle force, we investigated mechanical energetics of the rectus femoris in 7 individuals who experienced rapid, transient unilateral belt accelerations during walking. We hypothesized that: H1) the rectus femoris would actively lengthen on the perturbed leg during the perturbed stride and H2) on the contralateral leg the rectus femoris would reflect the mechanical energetic demand at the knee and leg levels. H1 was partially supported, with the rectus femoris fascicles being decoupled from muscle-tendon unit lengthening. H2 was not supported, with the rectus femoris best reflecting the energetic role of the hip, as opposed to the knee or leg. Overall, these findings provide a first estimate of the variety of roles proximal muscles play in maintaining stability and lay the groundwork for additional *in-vivo* measurement informed multi-scale analyses of perturbed locomotion.

## Introduction

As animals move through their environments, they exhibit an impressive ability to maintain stability despite external mechanical disturbances (*e*.*g*., rocks, sidewalks, ice). The high-level goal of maintaining a locomotor trajectory is accomplished through stabilizing responses mediated by 1) feedforward strategies (*e*.*g*., foot placement (1,2)), 2) feedback mechanisms (*e*.*g*., reflexes (3,4)]), and 3) intrinsic mechanical properties of biological structures (*e*.*g*., muscle-tendon interactions (5–7)). While stabilizing mechanical responses have been studied at different levels of musculoskeletal description (*e*.*g*., center of mass [COM], legs, joints, muscles), relating responses *across* different levels remains difficult. To address this challenge, a paradigm was recently developed which used mechanical energy to both quantify the explicit energetic demand of a perturbation and to serve as a “common currency” with which to relate responses across different levels of musculoskeletal description in humans (8).

This approach leveraged the idea that during level split-belt treadmill walking, if both belts are moving at the same constant speed, both the net work performed by each leg on the COM and the corresponding treadmill belt must be zero over a stride on average, so the net work performed by the overall leg (*i*.*e*., the sum of both COM and treadmill contributions), must be zero as well (9,10). Barring modeling inaccuracies, this overall leg work should be equal to the sum of the work of all lower limb joints (11). In (8), transient increases in unilateral belt speed were targeted to either early or late stance to perturb the flow of mechanical energy. In this non-steady state condition, net mechanical work was performed by each leg on the COM and the corresponding treadmill belt. The difference between these non-steady state overall leg and joint level mechanical work contributions and their corresponding steady state values constituted the mechanical energetic demand of the perturbation. By relating changes in joint and overall leg work, this approach allowed for investigation of which joints best reflected leg level demands. For the perturbed leg, on both the perturbed and first recovery stride, changes in ankle (vs. knee or hip) work best reflected changes in overall leg work, in agreement with literature from speed and ground height changes (12–16). However, on the contralateral leg, the knee strongly reflected overall leg level demands, spanning both net positive and negative work during the perturbed stride. This finding suggested proximal joints and muscles of the leg may play an important role in mediating transient energetic demands.

The goal of this work is to extend the perturbation energetics paradigm and investigate the role of proximal leg musculature in responding to rapid unilateral belt accelerations. As there is currently no gold-standard approach to quantifying human *in-vivo* fascicle lengths of proximal muscles during dynamic tasks, to accomplish our overarching goal we developed a custom automated application which leveraged deep-learned models for aponeurosis detection in conjunction with fascicle detection based on other recent fascicle tracking algorithms (17,18) to estimate fascicle lengths from B-mode ultrasound images. These fascicle lengths were used together with *in vivo* muscle activations to estimate fascicle and MTU forces (19), in addition to the respective mechanical powers. Of the quadriceps and hamstring muscles, we focused our analysis on the rectus femoris because 1) we sought to determine whether the extension of the perturbed limb by the perturbation would be apparent at the level of proximal muscle fascicles, as opposed to being buffered by series elasticity (7,20), and 2) an increased knee extensor moment is a major feature of the recovery strategy on the contralateral leg following a belt acceleration during walking (21). Since proximal muscles are considered to have less compliant tendons relative to distal muscles (22,23), our first hypothesis (H1) was that the rectus femoris would actively lengthen on the perturbed leg during the perturbed stride, resulting in increased negative fascicle and MTU power and work relative to steady state. On the contralateral leg, because the knee reflected overall leg level energetic demands (8), and the functional behavior of the rectus femoris fibers and MTU (predominantly damper-like; (24)) is more similar to the knee than the hip (predominantly motor-like; (15)), our second hypothesis was that the rectus femoris would reflect contralateral knee energetic demands and therefore also overall leg energetic demands.

## Methods

### Experimental Protocol

Data were collected from 7 healthy able-bodied individuals (5 males, 2 females, mean [SD]: 25 [2] years, 178.5 [12.1] cm stature, 72.7 [13.3] kg) as part of a study investigating leg and joint level energetics (8). Participants walked on an instrumented split-belt treadmill (CAREN, Motek, Netherlands), and after 5 minutes of acclimation at 1.25 m s^-1^ (25), experienced rapid (15 m s^-2^ acceleration), transient (∼340 ms duration, ∼33% of the perturbed gait cycle) increases in unilateral belt speed (1.25 m s^-1^ to 2.5 m s^-1^) targeted to either early or late stance (Figure 1). The algorithm by which these perturbations were delivered is described elsewhere (26). Each participant experienced 40 total perturbations in a randomized order (both legs, early and late stance onset timings, and 10 repetitions of each leg/timing combination), with 30-40 steps between perturbations to allow for return to steady state walking (27). All participants (recruitment period: 8/28/2020 - 3/6/2021) provided written informed consent and all protocols were approved by the Institutional Review Board at the Georgia Institute of Technology (Protocol H20163). The sample size was determined based on pilot data indicating a sample of 6 participants would be capable of detecting a significant (p < 0.05) effect of speed on average fascicle length over a stride in proximal leg muscles with 82% power.

**Figure 1.**
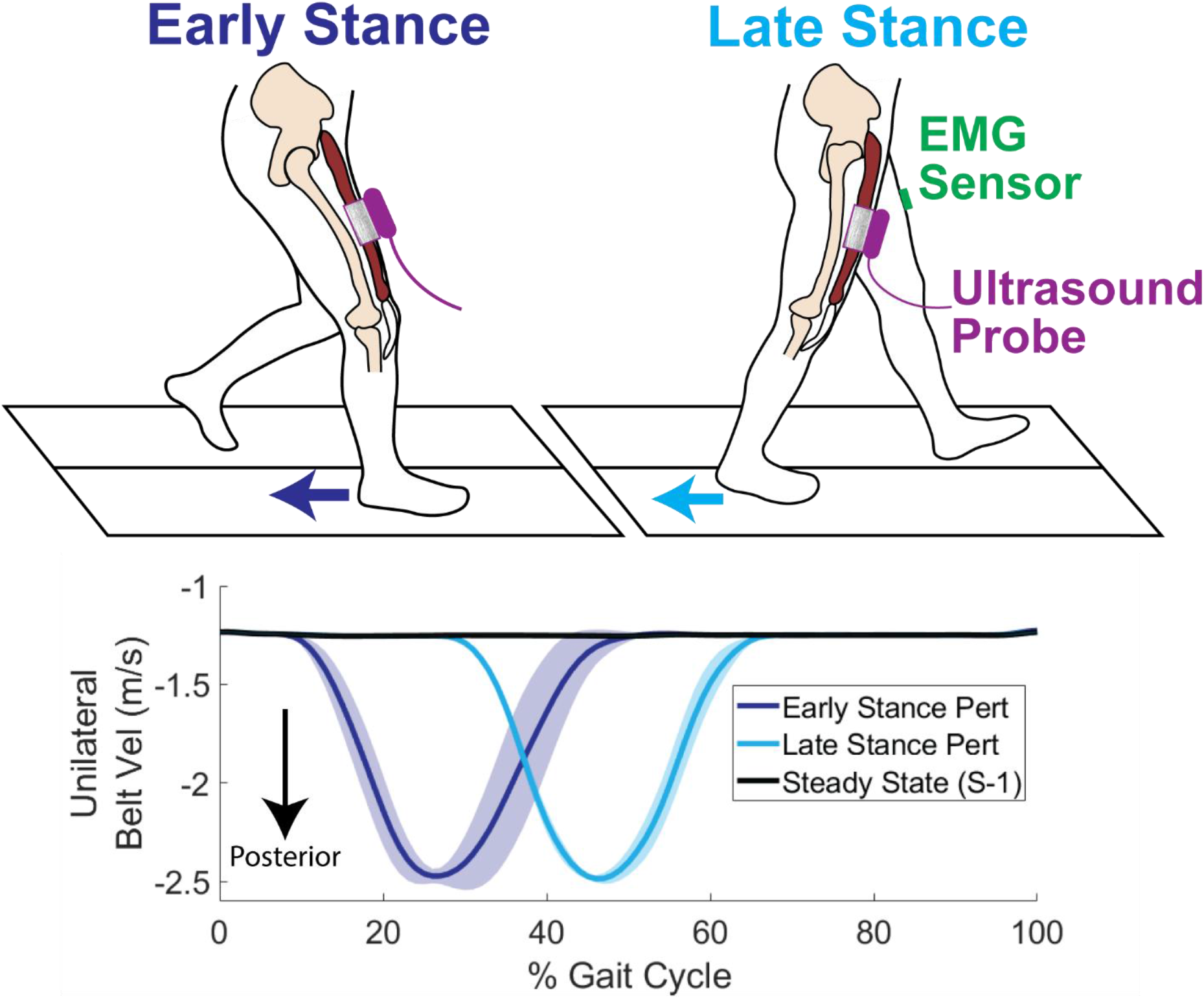
Conceptual overview of the treadmill perturbation and muscle-level measurements. Unilateral treadmill belt velocity traces show across-subject ensemble averages. Shaded regions represent ±1 standard deviation.

### External Kinematics and Kinetics

Ground reaction forces were collected at 2000 Hz from an instrumented split-belt treadmill (CAREN, Motek, Netherlands). A full-body marker set consisting of 67 retroreflective markers (modified Human Body Model 2; (28)]) was used to track bony landmarks and body segments (head, hands, forearms, upper arms, torso, pelvis, thighs, shanks, and feet). Marker trajectories were collected at 200 Hz using a 10-camera motion capture system (Vicon; Oxford, UK). For each participant, a generic full-body musculoskeletal model (22 segments, 37 degrees of freedom; (29)) was scaled in OpenSim 4.0 (30) using a static standing trial to generate a participant-specific model. The metatarsophalangeal and subtalar joints were locked to treat the foot as a rigid body. All trials were manually inspected to ensure crossover steps were removed from analysis, leaving 222 successful trials. Joint angles were calculated from marker trajectories using the OpenSim Inverse Kinematics tool. Joint moments were calculated with the OpenSim Inverse Dynamics tool using both joint angles and ground reaction forces applied to the calcanei of the scaled models. Time-varying moment arms of the rectus femoris to the hip and knee joints in addition to MTU lengths and velocities were calculated using the OpenSim Muscle Analysis tool using joint angles and the scaled models. Kinematic and kinetics were low-pass filtered using 4^th^ order zero-phase Butterworth filters with cutoff frequencies of 6 and 15 Hz, respectively. Strides were segmented using a 30 N threshold applied to vertical ground reaction force.

### Leg and Joint Energetics

Joint mechanical powers were calculated as the product of joint angular velocities (obtained by differentiating joint angles with respect to time) and joint moments. Mechanical powers of each leg were calculated using a modified individual limbs method (11,31) as the sum of 1) power flowing from each leg to the COM, 2) power flowing from each leg to the treadmill, and 3) peripheral power of the leg segments moving relative to the COM of the whole body. Leg to COM power was calculated as the dot product of the corresponding ground reaction force and COM velocity. COM velocity was calculated as the time derivative of COM position from the Body Kinematics tool in OpenSim using both joint angles and scaled models. Leg to treadmill power was calculated as the product of anteroposterior leg force (equal and opposite to ground reaction force) and treadmill velocity (collected at approximately 70 Hz). Peripheral leg power was quantified by adding together the time derivatives of the rotational and translational components of kinetic energy of the leg segments (11,32,33) as calculated from inertial estimates from the scaled models and segment velocities from the OpenSim Body Kinematics tool.

### Electromyography

Surface electromyography (EMG) was collected at 2000 Hz (Avanti, Delsys, Natick, MA) from 7 lower limb muscles of the left leg: the rectus femoris (RF), tibialis anterior, soleus, medial gastrocnemius, vastus medialis, biceps femoris, and gluteus maximus. EMG sensors were placed on the bellies of each muscle following light abrasion of the skin surface. All EMG envelopes were calculated by first high-pass filtering EMG signals using a 10 Hz zero-phase Butterworth filter, then full-wave rectifying the signal, then low-pass filtering the signal with a 30 Hz zero-phase Butterworth filter (34,35). Signal offsets were first removed by subtracting the minimum envelope value during a relaxed static trial. These demeaned envelopes were then normalized by the difference between the maximum within-participant envelope value across all perturbations and the minimum resting value, similar to previous work (34,36). To estimate muscle activations, this normalized signal was then passed through first-order dynamics (37) with an activation time constant of 35 ms, and a deactivation time constant of 58 ms, corresponding to a mixed fiber type muscle (19), which is applicable to the RF (38). Since the RF is the primary muscle of interest in this work, activations of other muscles are included as a supplementary figure (Figure S1).

### Ultrasonography

B-mode cine ultrasound images were collected at approximately 115 Hz from the RF of the right leg using an ArtUS EXT-1H acquisition unit and a 60 mm long LV8-5N60-A2 probe (Telemed, Vilnius, LT). The probe was placed over the muscle belly at a consistent longitudinal location to the RF EMG sensor on the left leg. The probe was aligned such that fascicles and both superficial and deep aponeuroses could be clearly visualized. The imaging angle of the probe was also varied in EchoWave software (Telemed, Vilnius, LT) to provide the best contrast of the fascicles and aponeuroses – between 0 and −10 degrees based on the participant. An external trigger was used to synchronize motion capture and ultrasound image timestamps.

### Custom Ultrasound Tracking Application

A custom application was developed in Matlab R2019b (Mathworks, Natick, MA, USA) to estimate proximal muscle fascicle lengths and pennation angles from B-mode ultrasound images during dynamic tasks. The application consists of two parts. <u>Part 1</u> involves training study-specific deep-learned models for detection of superficial and deep aponeuroses. <u>Part 2</u> uses the identified aponeurosis locations to define a region of interest and estimates representative fascicle parameters from hyperechoic “snippets” of fascicles within that region. The application workflow is shown in Figure 1B.

### Part 1 – Aponeurosis Identification using Deep-Learning

Part 1 of the application consists of 3 steps. <u>First</u>, an image set is generated to train and validate deep-learning models for identification of superficial and deep aponeuroses. The overarching design strategy of this approach is to “overfit” models to perform best on the remaining images of a study. This approach allows for rapid training of models with a small dataset on a mid-range desktop computer and avoids having to collect and manually track a data set that generalizes across tasks, probes, participants, etc. (39). The drawback of this approach is that the trained models are likely study specific, limiting reproducibility. In this study, a set of 490 images was used for training and validating models. This image set consisted of 35 frames equally spaced in time across 1 perturbed gait cycle from both the ipsilateral and contralateral legs for all 7 participants. From the identified set of images, the superficial and deep aponeuroses are then manually traced to produce binary masks (roipoly, Matlab). The <u>second</u> step is training separate models for identification of pixels belonging to either superficial or deep aponeuroses. Both the raw images and masks are resized to 512x512 pixels for use in model training. A U-net model architecture is used based on previous success in identifying pixels corresponding to fascicles and aponeuroses (40,41). For both the superficial and deep aponeurosis models, the total image set is first randomized, then split into training (80% of the image set) and validation sets (20% of the image set). In this study, each model took approximately 1 hour to train (trainNetwork, Matlab) on a desktop computer with an Nvidia GTX 1660 Ti graphics card. From the validation set, pixel-wise mean classification accuracies were 96% and 94% and mean intersection over union values were 94% and 91% for the superficial and deep aponeuroses, respectively. The <u>third</u> step is applying the trained models to the broader dataset (∼111,000 frames) and generation of first order polynomials to represent the superficial and deep aponeuroses which defines the region of interest for fascicle detection. This is accomplished by generating masks of both the superficial and deep aponeuroses from each frame of ultrasound data using the trained models (semanticseg, Matlab), identifying the largest object by area in each of the predicted masks, and then fitting a first order polynomial based on the centroid and orientation of each identified object (regionprops, Matlab). The intersection points of these superficial and deep first order polynomials with the proximal and distal borders of the image serve as the vertices bounding the region of interest. These regions of interest are then used in Part 2.

### Part 2 – Representative Fascicle Estimation

A GUI-based Matlab application was developed to allow for streamlined visualization of representative fascicle measurements and correction of aponeurosis fits. The functionality of this application is separated into 4 steps. <u>First</u>, the raw images, with fascicles running from top left to bottom right, are filtered using adaptive histogram equalization (adapthisteq, Matlab) to improve contrast of hyperechoic structures. To account for angled images, the user is prompted to enter the image angle, which is used to first shear the image to have vertical edges, then applies adaptive histogram equalization (adapthisteq, Matlab), then returns the image to the original angle. This step is necessary as the border outside an angled image introduces artifacts during filtering. From the filtered image, the area between the aponeuroses is identified as the region of interest, with all structures outside of that area being removed. The region of interest is binarized (imbinarize, Matlab) using a Frangi-type vessel enhancement filter (fibermetric, Matlab; thickness = 7, sensitivity = 100) as has been used in previous tracking algorithms (18,42,43). From this binarized image, the orientations and major axis lengths of each separate structure are computed (regionprops, Matlab). The user also has the option of further limiting the region of the image passed through to fascicle detection by applying an additional manual mask to these results, which may be warranted if the muscle of interest has compartments of different architectures. The <u>second step</u> is identification of the “snippets” in the binarized image which can be attributed to fascicles. Similar to (17), only structures with major axis lengths greater than 4 mm, area to length ratios less than 12, and with orientations relative to the deep aponeurosis within specified bounds are used to estimate the representative fascicle of each frame. The range of acceptable pennation angles relative to the deep aponeurosis was set between 6 and 20 degrees, based on *in-vivo* and cadaveric measurements of RF architecture (44,45). <u>The third step</u> is assessment of the thickness of the muscle. This is calculated as the length of a line segment running perpendicular to the deep aponeurosis which intersects both the superficial aponeurosis and the midpoint of the deep aponeurosis. <u>The fourth step</u> is calculating representative fascicle parameters. The representative pennation angle of a frame is calculated as the circular mean weighted by snippet length from all accepted snippets, as adapted from (17). The representative fascicle length is then calculated as the thickness divided by the sine of this representative pennation angle (18,43,44). Following completed processing of a trial, the ensemble averaged and time normalized fascicle lengths and pennation angles are visualized. The application interface during use is included as a supplement (Figure S2). For all subsequent analyses, fascicle lengths and pennation angles were low-pass filtered at 6 Hz with a zero-phase 4^th^ order Butterworth filter. Fascicle velocities were calculated as the time derivative of filtered fascicle lengths.

### Estimation of Fascicle and Muscle-Tendon Unit Energetics

Fascicle force was estimated from a Hill-type model of muscle force generation driven by a combination of experimentally measured fascicle lengths and muscle activations similar to (19):

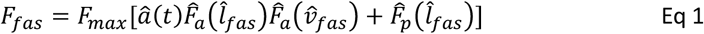

Where *F*_*max*_ was maximum isometric fascicle force, â *(t)* was estimated muscle activation from 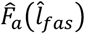 EMG, was the normalized relationship between active force and fascicle length,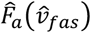 was the normalized relationship between active force and fascicle velocity, and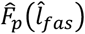 was the relationship between passive force and normalized fascicle length. 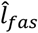and 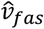 were fascicle length and velocity, respectively, normalized to fascicle slack length, *l*_*0,fas*_ (mean [s.d.]: 145.5 [22.4] mm across all participants). *l*_*0,fas*_ was determined as the length when force began to appear in the tendon, which based on previous measurements of patellar tendon force in humans (46) was approximated for each participant to be the average fascicle length at 50% of the walking gait cycle at 1.25 m s^-1^. MTU force was calculated as:

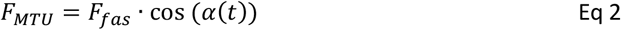

Where *α(t)* is pennation angle. Maximum fascicle force was calculated as:

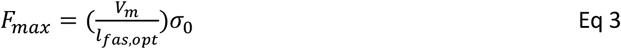

Where *V*_*m*_ is muscle volume estimated from each participants height and mass (47), σ_0_ is specific muscle stress (48,49), and *l*_*fas,opt*_ is optimal RF fiber length estimated from scaled OpenSim models of each participant (50). From (51,52), the normalized active force-length relationship was calculated as:

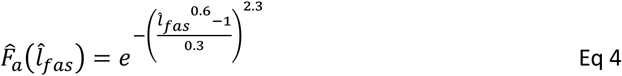

For normalized fascicle lengths greater than unity, the normalized passive force-length relationship (53) was calculated as:

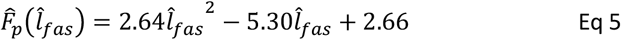

For normalized fascicle lengths less than or equal to unity, the normalized passive force was set to 0. For shortening fascicles 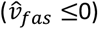 the normalized force-velocity relationship was given by (52):

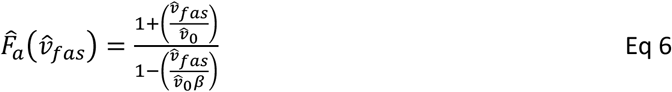

For lengthening fascicles 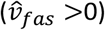 the normalized force-velocity relationship was given by:

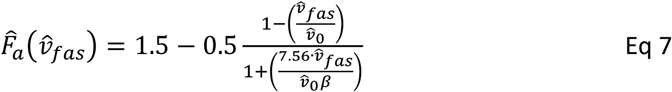

Where *β* was the fiber type dependent curvature of the force-velocity relationship (52). Since the human RF is of a mixed fiber type (38), *β* was determined to be the average of the constants determined from literature of both slow and fast fiber types (0.18 and 0.29, respectively; (19)). Similarly, 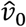, the maximum intrinsic speed of the fascicles, was estimated as the average (7.5 s^-1^) of values from slow and fast fibers (5 and 10 s^-1^, respectively (54,55)).

Since EMG and ultrasound images were collected from different legs, for all fascicle or MTU force, power, and work calculations information from both legs was combined. This was achieved by ensemble averaging activations, fascicle length/velocities, and MTU lengths/velocities, and fascicle pennation angles across all iterations of a given perturbation timing and side within each participant. Thus, within-participant and within-condition ensemble averaged timeseries were calculated for each stride (*e*.*g*., one trace for all RF activations ipsilateral to the perturbed leg for participant 1 during early stance perturbations). Using these combined traces, fascicle power was calculated as the product of fascicle force and fascicle velocity, and MTU power was calculated as the product of MTU force and MTU velocity, where fascicle or MTU lengthening resulted in negative power, respectively. See Figure 2 C for an overview of fascicle and MTU force and power calculations. Fascicle and MTU work values were calculated by integrating the ensemble averaged curves (trapz, Matlab) and then multiplying by the average stride duration of the traces divided by the number of normalized time points of each stride (101). To compare leg or joint level mechanical work values with fascicle and MTU mechanical work, mechanical work at the leg or joint levels was also averaged across all perturbation iterations within each participant.

**Figure 2.**
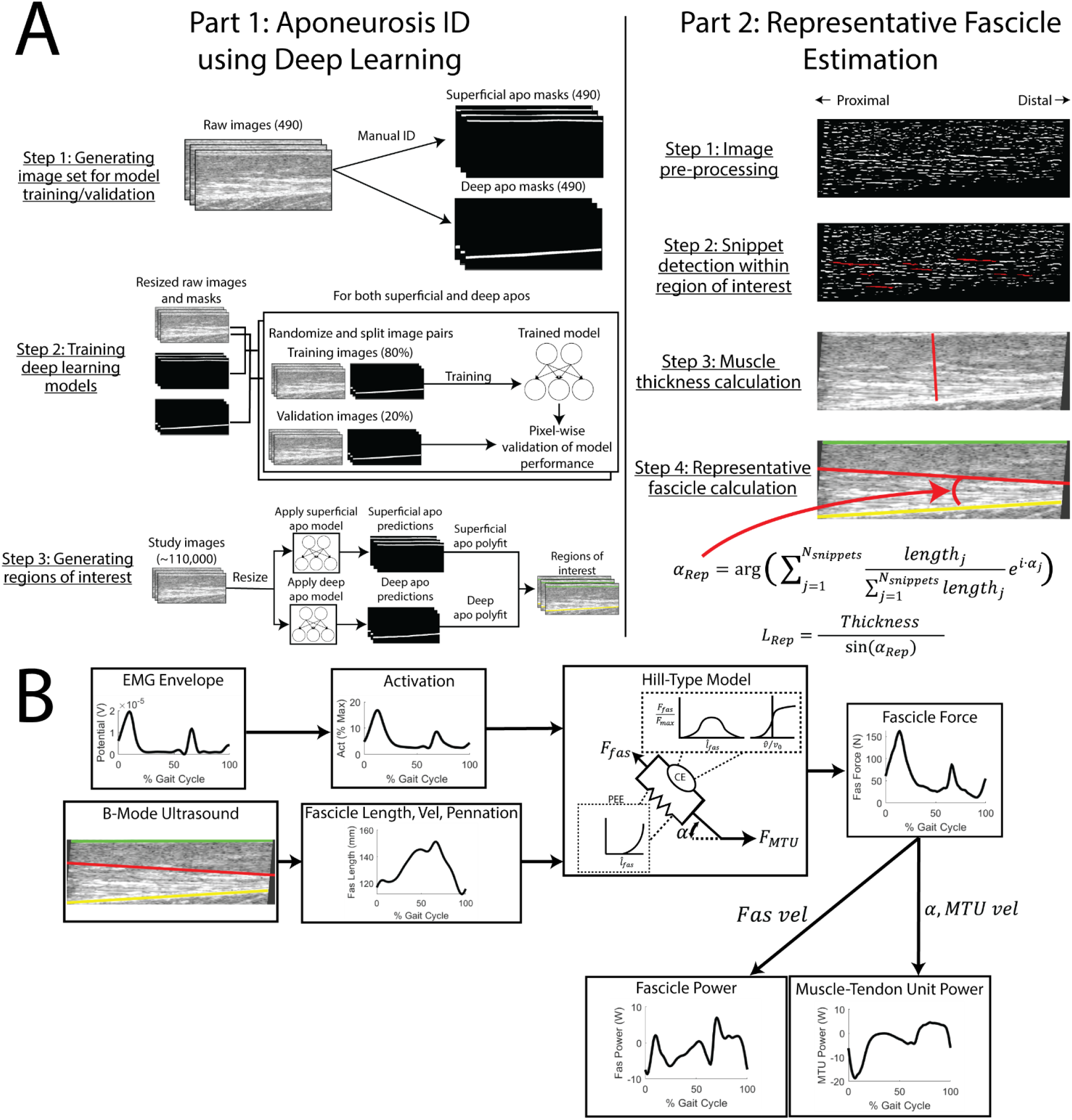
A) Overview of custom fascicle tracker application. α = fascicle pennation angle, L_Rep_ = representative fascicle length. B) Overview of fascicle and muscle-tendon unit energetics computation. CE = contractile element, PEE = passive elastic element.

### Statistical Analysis

Statistical analyses were performed in Matlab R2019b (Mathworks, Natick, MA, USA). To investigate differences in timeseries outcomes (e.g., fascicle lengths, activations, mechanical powers for H1), statistical parametric mapping (SPM; (56)) was used to run paired t-tests to compare early or late stance traces to steady state traces (*i*.*e*., the pre-perturbation stride) without needing to correct for multiple comparisons. While all statistical differences were reported, alterations in stride times due to the perturbations led to phase shifts in the gait-cycle normalized data that resulted in some periods of statistical significance that were not determined to be meaningful. Further, p-values greater than the significance threshold (*i*.*e*., 0.05) were approximated by solving p-value associated with the maximum t-statistic during the entire timeseries. This approach has not yet been validated (57) and in some cases resulted in values less than the significance threshold. Such cases are denoted with *ns*. All timeseries data were represented in relation to the gait cycle of the perturbed (i.e., ipsilateral) leg. For discrete outcome measures such as mechanical work, paired t-tests were used to compare early or late stance perturbation conditions to steady state levels (H2 and H3). To evaluate the functional role of the RF in relation to the hip, knee, and overall leg (H2), linear regressions were used to relate changes in fascicle or MTU work to changes in hip, knee, and overall leg work relative to steady state levels (i.e., the work of the perturbed stride – work of the previous unperturbed stride, S-1). Since the hip had 3 degrees of freedom in these analyses, hip work was calculated as the integrated sum of all 3 components to better account for overall leg work. Analyses were limited to the unperturbed, perturbed (S0), and first recovery strides (S+1) as a return to steady state in external kinetics and kinematics was largely achieved after the first recovery stride (8). Because data were collapsed across perturbation iterations, for each leg these regressions contained 14 points (7 participants x 2 timings) instead of 222 from all perturbations. Significance was concluded for p-values ≤ 0.05.

## Results

### During perturbations, rectus femoris fascicle rotation decouples length changes between MTU and fascicles levels, with increased activation driving active lengthening

For both early and late stance perturbations, during the perturbed stride (S0) on the ipsilateral leg there was no significant length change of RF fascicles (early: p = 0.679; late: p = 0.078), while there was significant lengthening of the MTU (early: p < 0.001; late: p = 0.022; Figure 3). This decoupling during perturbations appeared to be caused by increases in pennation angle (early: p = 0.080; late: p = 0.180) offsetting increases in muscle thickness (early: p = 0.201; late: p=0.033 *ns;* Figure 3). Further, although not statistically significant, an increase in RF activation relative to steady state levels was also observed (early: p = 0.080; late: p = 0.036 *ns*; Figure 4). The natural lengthening of RF fascicles from mid to late stance coupled with increased activation resulted in increased fascicle forces during mid to late stance for early stance perturbations and around toe-off for late stance perturbations (p<0.001 for both; Figure 4). This increased force during lengthening resulted in larger negative powers and work at the level of the RF fascicles and the MTU (Figure 5). At the level of fascicles, negative powers during stance were not significantly different from steady state for either timing (early: p = 0.025 *ns*; late: p= 0.475), while significantly more negative work was performed over the perturbed stride for both timings (early: p < 0.001; late: p = 0.039). At the level of the MTU, there were significantly larger negative power relative to steady state for both timings (p < 0.001 for both), which also resulted in significantly more negative work (early: p = 0.002; late: p = 0.003).

**Figure 3.**
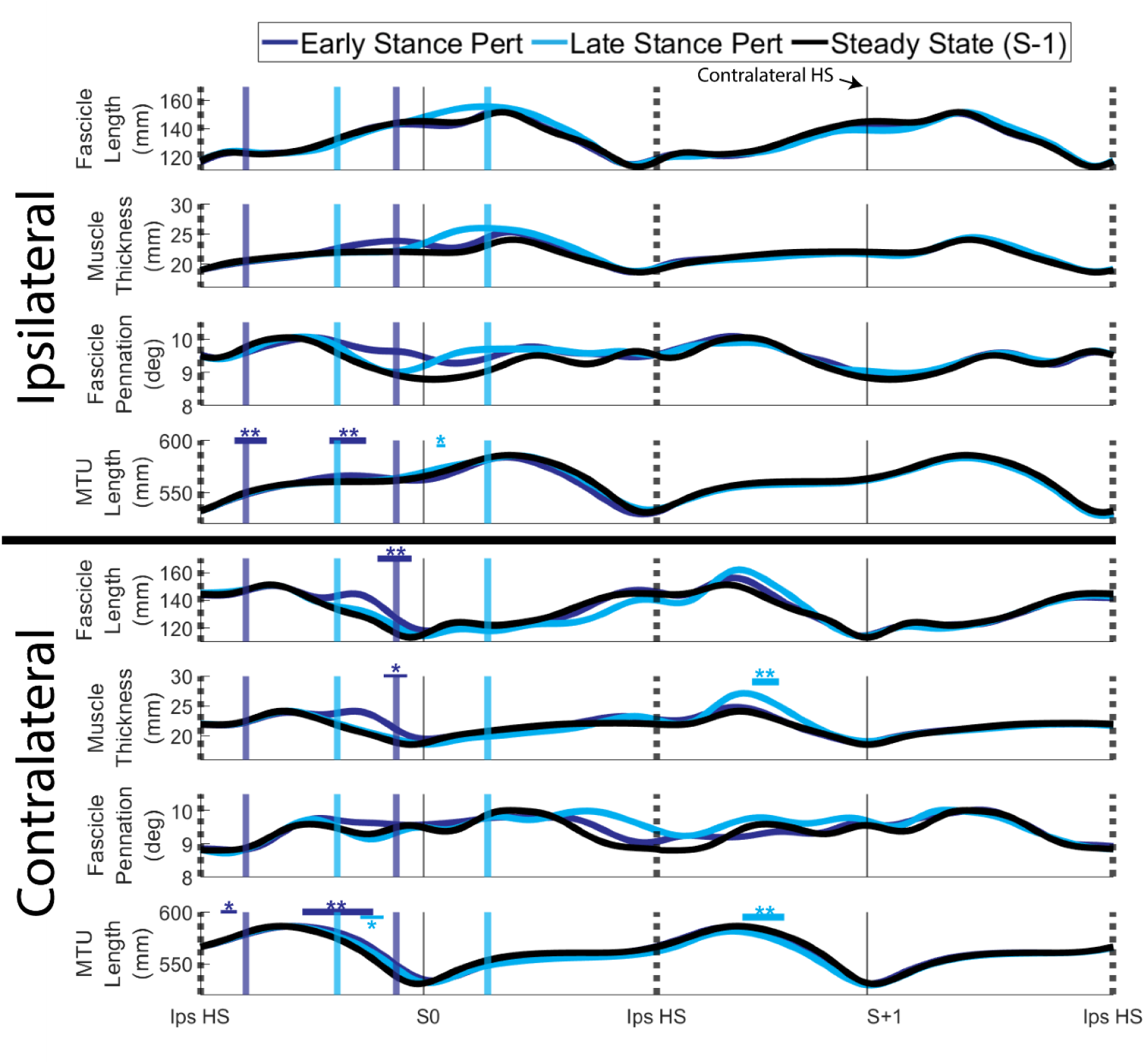
Rectus femoris fascicle and MTU dynamics on both legs relative to the perturbation across all participants (n=7, 222 perturbations), time-normalized to gait cycles. Standard deviations were omitted for clarity. Instances when curves significantly deviated from steady state (S-1) as determined by statistical parametric mapping two-tailed t-tests are identified with thick horizontal lines and ** for p < 0.001, and thin horizontal lines and * for p < 0.05. Solid colored vertical lines indicate average start and end times of the perturbations. Dotted/solid black vertical lines indicate instances of ipsilateral/contralateral heelstrike, respectively. “Steady State” strides were the strides preceding the perturbed stride (S-1).

**Figure 4.**
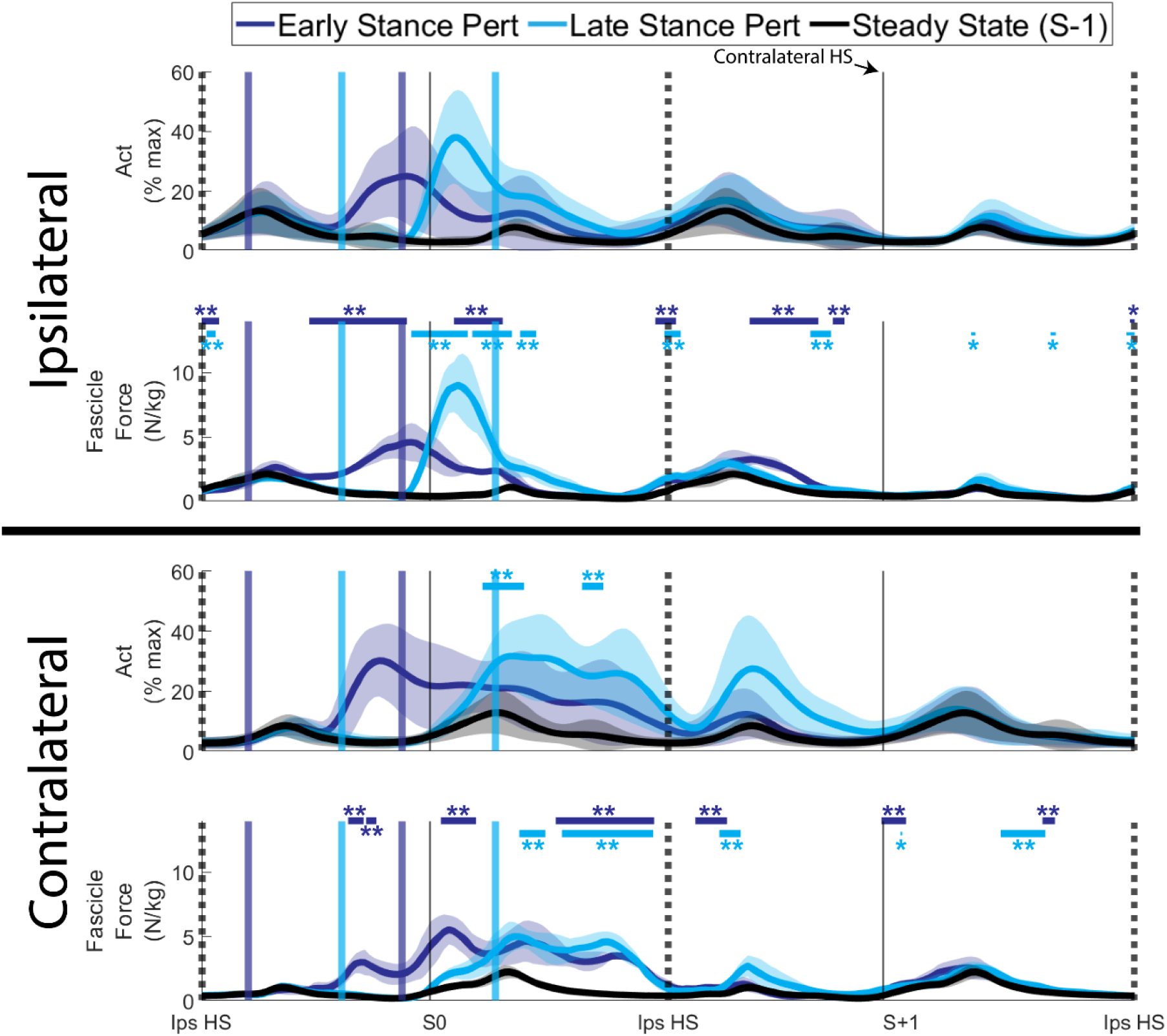
Rectus femoris activations (% max across all trials) and fascicle forces on both legs relative to the perturbation across all participants (n=7, 222 perturbations), time-normalized to gait cycles. Shaded regions represent ±1 standard deviation. Instances when curves significantly deviated from steady state (S-1) as determined by statistical parametric mapping two-tailed t-tests are identified with thick horizontal lines and ** for p < 0.001, and thin horizontal lines and * for p < 0.05. Solid colored vertical lines indicate average start and end times of the perturbations. Dotted/solid black vertical lines indicate instances of ipsilateral/contralateral heelstrike, respectively. “Steady State” strides were the strides preceding the perturbed stride (S-1).

**Figure 5.**
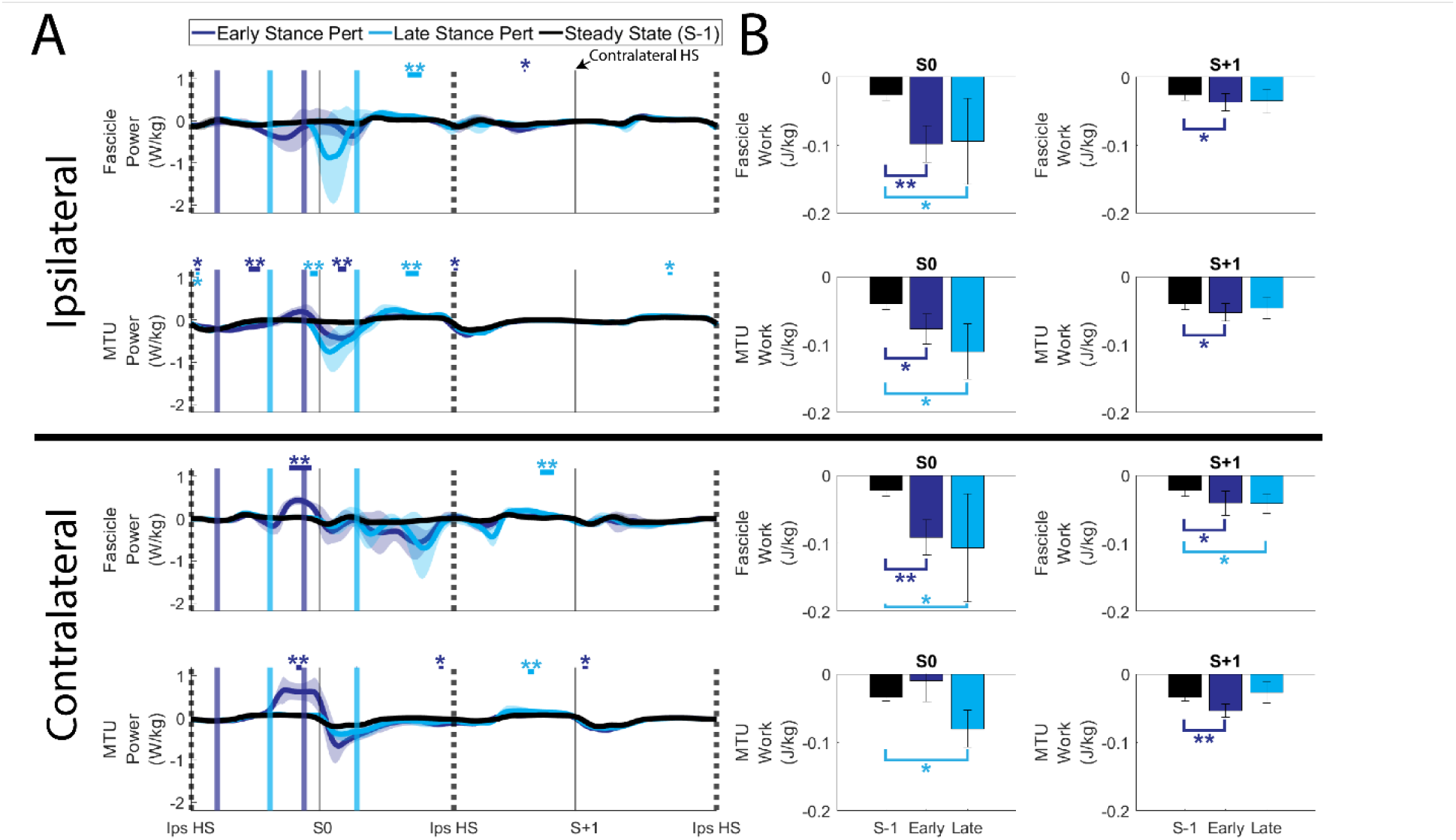
A) Rectus femoris fascicle and MTU mechanical powers on both legs relative to the perturbation across all participants (n=7, 222 perturbations), time-normalized to gait cycles. Shaded regions represent ±1 standard deviation. Instances when curves significantly deviated from steady state (S-1) as determined by statistical parametric mapping two-tailed t-tests are identified with thick horizontal lines and ** for p < 0.001, and thin horizontal lines and * for p < 0.05. Solid colored vertical lines indicate average start and end times of the perturbations. Dotted/solid black vertical lines indicate instances of ipsilateral/contralateral heelstrike, respectively. “Steady State” strides were the strides preceding the perturbed stride (S-1). B) Rectus femoris fascicle and MTU work over the perturbed (S0) and first recovery stride (S+1) on both sides relative to the perturbation. Within-participant paired two-tailed t-test results comparing perturbed to steady state (S-1) values are represented as ** for p<0.001 and * for p<0.05. The rectus femoris MTU best reflects the role of the contralateral hip

### Rectus femoris mechanical energetics on the contralateral leg and first recovery stride

On the contralateral leg to the perturbation, during the perturbed stride (S0), early stance perturbations were associated with longer fascicle and MTU lengths relative to steady state, followed by shortening (p < 0.001) to allow lengths to return to steady state levels around contralateral heelstrike (Figure 3). This shortening coincided with increased RF activation (p = 0.025 *ns*) to result in increased forces (p < 0.001; Figure 4) and larger positive fascicle and MTU powers (p < 0.001; Figure 5). However, for fascicles, these larger positive powers were offset by larger negative powers later in the gait cycle which resulted in fascicles performing larger amounts of negative work over the perturbed stride relative to steady state (p < 0.001; Figure 5). Conversely, for the MTU, the larger negative power during the perturbation was not fully offset and the net work performed by the MTU was greater than steady state levels, though this difference was not significant (p = 0.076; Figure 5).

For late stance perturbations, changes from steady state were predominantly limited to increased activations (p < 0.001) and forces (p < 0.001) coinciding with steady state levels of fascicle and MTU lengthening. This resulted in larger negative fascicle (p = 0.018) and MTU works (p = 0.002) relative to steady state.

Across both legs, the largest deviations from steady state on the first recovery stride (S+1) occurred following late stance perturbations on the contralateral side. While contralateral MTU lengths were shorter relative to steady state (p < 0.001), the contralateral RF fascicles lengthened then shortened during swing, although not significantly (p = 0.082), due to increasing pennation offsetting increased thickness (p < 0.001; Figure 3). Increased RF activations drove higher forces (p < 0.001; Figure 4), which resulted in a period of negative followed by positive fascicle power, which on net over the first recovery stride led to slightly more negative work being performed by the fascicles relative to steady state (p = 0.018; Figure 5). In general, changes in both ipsilateral and contralateral fascicle and MTU work from steady state levels were minimal during the first recovery stride.

Changes in RF MTU work most strongly related to changes in hip work on the leg contralateral to the perturbation during both the perturbed and first recovery strides. Fascicles had much weaker relationships to joint level demands compared to MTUs (max R^2^ = 0.22, p > 0.051), although the direction of the relationships was general consistent across fascicle and MTU levels for a given leg and stride (Figure 6). Further, relationships between both fascicle and MTU levels with leg level demands were consistently weak (max R^2^ across all = 0.22, p > 0.051)

**Figure 6.**
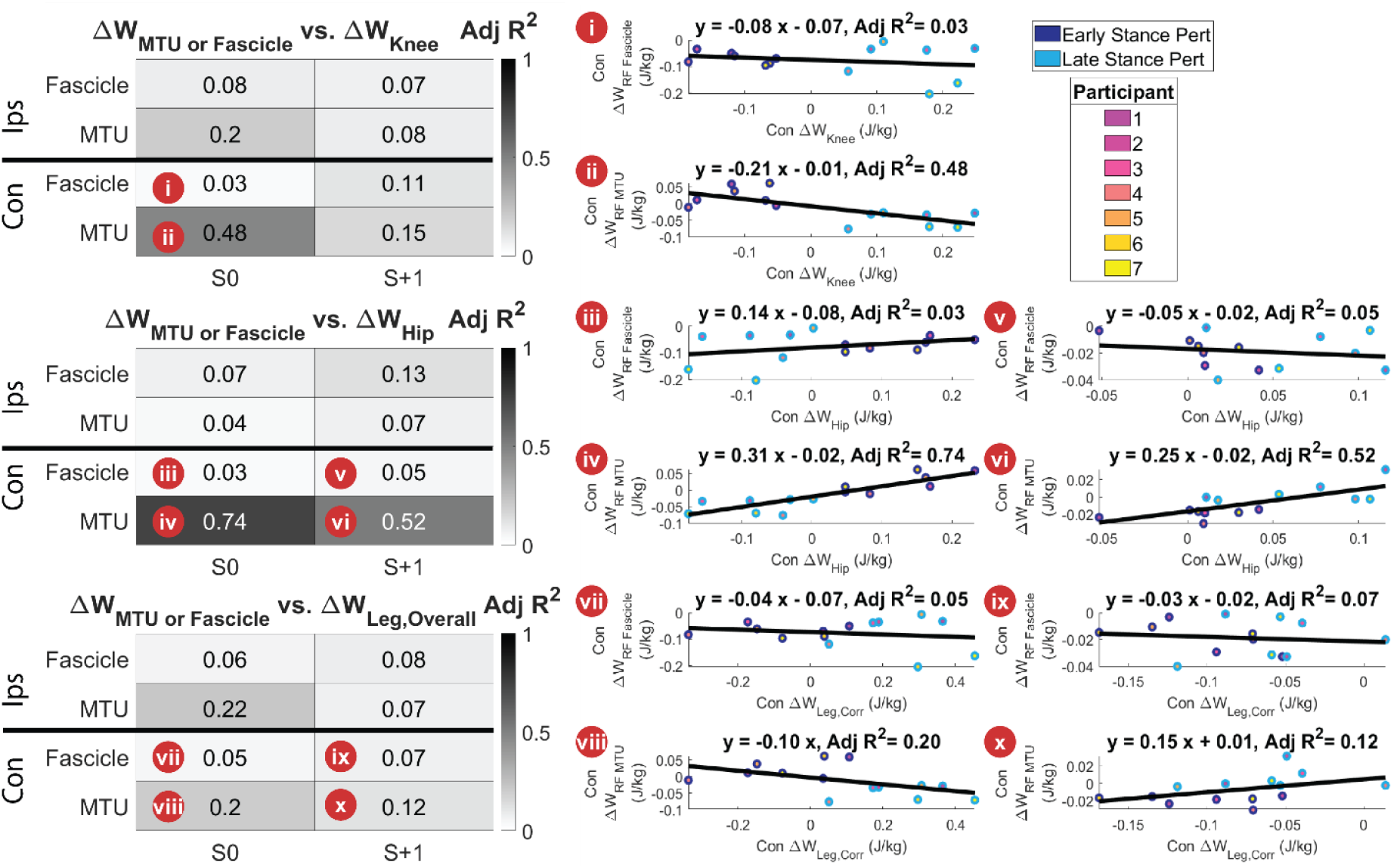
Relationships between rectus femoris fascicle or MTU work and the knee, hip, and overall leg. Heatmaps show adjusted R^2^ values for linear regressions between differences in fascicle or MTU work over a stride from steady state, and differences in knee, hip, or corrected leg work over a stride from steady state (S-1). Note that fascicle and MTU data were collapsed across iterations and sides, resulting in 14 data points (n=7, 2 perturbation timings) per scatter from the original 222 perturbations. (i-x) Scatter plots for selected fascicle and MTU relationships of interest.

On the perturbed stride (S0), contrary to our hypothesis, the RF MTU had the strongest positive relationship with the hip (MTU R^2^ = 0.74, p < 0.001) and a weaker negative relationship with the knee (MTU R^2^ = 0.48, p = 0.004). On the first recovery stride (S+1), there was also a strong positive relationship between RF MTU and hip demands (R^2^ = 0.52, p = 0.002).

## Discussion

The main objective of this work was to evaluate the role of the RF in mediating transient mechanical energy demands during perturbed walking in humans. Using unilateral belt accelerations on a split-belt treadmill, we imposed quantifiable mechanical energy demands on both legs during walking and extended our previous analysis relating joint and leg level demands down to the level of the RF MTU and fascicles (8). To estimate mechanical power and work of the RF for this approach, we used *in-vivo* RF fascicle lengths measured using B-mode ultrasound and a custom fascicle tracking application in conjunction with EMG to drive a Hill-type model of force generation. Our first hypothesis (H1) sought to address the energetic implications of the perturbation itself – specifically, we hypothesized that on the perturbed leg, the perturbation would result in negative RF power and work because of active muscle lengthening. Our results partially supported this first hypothesis, with the kinematics of the perturbation resulting in lengthening of the RF MTU relative to steady state walking, but not the RF fascicles. This decoupling was due to an increase in fascicle pennation angle offsetting a potential increase in fascicle length due to increased muscle thickness. While decoupling during eccentric contractions due to changes in architectural gearing have previously been observed, such changes are due to a decrease in thickness being offset by a decrease in pennation angle. This discrepancy is likely due to changes in slope of the deep aponeurosis – during perturbations we observed that the deep aponeurosis exhibited an increased slope relative to steady state, thereby increasing the measured thickness and increasing the measured pennation angle (Figure S3). Linear regressions between deep aponeurosis angle and pennation angle (R^2^=0.37, p<0.001) in addition to the sine of deep aponeurosis angle and thickness (R^2^=0.14, p<0.001) over the stance phase on the perturbed leg further support this relationship. However, our hypothesis was still partially supported since increased RF activation during the perturbation together with similar lengthening observed during steady state walking precipitated increased negative power and work. While this represents the first *in-vivo* estimate of rectus femoris mechanical energetics during perturbed walking, the dissipation induced by the perturbation at proximal MTU and fascicle levels is mechanically analogous to other transient, dissipation-inducing perturbations delivered at distal sites. Specifically, unexpected changes in substrate height have been extensively investigated in both hopping humans and running guinea fowl (5–7), with drops in ground height resulting in active lengthening of the plantarflexors. In perturbed human hopping, a drop in terrain height has been shown to extend the aerial phase and thus shift the onset timing of plantarflexor activation to be earlier relative to ground contact. This earlier activation in concert with coactivation of the tibialis anterior has been demonstrated to: 1) allow plantarflexor muscles to pre-emptively shorten before ground contact, allowing for lower, potentially less damaging muscle strains, 2) stiffen and stabilize the ankle, and 3) bias the muscle towards achieving the task demand (*i*.*e*., dissipation of injected energy) due to alignment of activation and fascicle lengthening (7,58). We now address each of those previous findings in light of our results. In our perturbation context, instead of delaying the onset of ground contact, we applied both perturbation timings within stance phase, which did not allow for a pre-emptive shortening during an unloaded period. However, even without this pre-emptive shortening, we observed steady-state magnitudes of fascicle lengthening during the perturbations. While buffering of length changes between the MTU and fascicle levels in other quadriceps muscles has previously been attributed to series elasticity (59), we found that such lengthening may be buffered by shape change of the deep aponeurosis. Consistent with previous work investigating changes in activation of leg musculature during walking on uneven terrain (60), we observed coactivation of muscles across the lower limb on the perturbed leg (Figure S1), with increases in plantarflexor activation occurring first and being the most prominent. However, a previous study specifically investigating unilateral treadmill belt accelerations did not see increases in tibialis anterior activity following plantarflexor activation, which could be attributed to the less intense perturbations not eliciting a whole-leg coactivation strategy (61). Lastly, on the perturbed leg, we previously found the overall leg level demand of early stance perturbations was generation of net work over a stride, and the demand of late stance perturbations was net zero work over a stride, since net positive work performed by the leg on the treadmill is offset by net negative work performed by the leg on the COM (8). Therefore, in contrast to tasks where plantarflexors have been shown to modulate their mechanical role to match that of the limb (5–7), during a treadmill acceleration task the RF on the perturbed leg predominantly serves to dissipate energy and thus does not directly represent the energetic role of the leg, knee, or hip (Figure 6).

Despite the RF not reflecting the role of joints or the leg on the perturbed side, since we previously identified the contralateral knee as reflecting the energetic demand on the contralateral leg, we hypothesized due to the similar “damping” functions of the knee and RF (H2) that the contralateral RF may reflect knee, and therefore leg level demands on the perturbed stride. In contrast to our hypothesis, the RF, particularly at the MTU (vs. fascicle) level, best reflected the role of the contralateral hip, while the relationship between the RF and the knee and leg were weaker and negative (Figure 6) This relationship was due to 1) early stance perturbations eliciting positive net work at the hip joint and the MTU and fascicles actively shortening during swing to produce a period of positive power and 2) late stance perturbations eliciting negative work at the hip joint and the MTU and fascicles actively lengthening during stance to produce almost exclusively negative power. In contrast, the work demands on the knee are opposite to that of the hip during these perturbation timings (*i*.*e*., net negative knee work during early stance perturbations, net positive knee work during late stance perturbations). These findings indicate that while the role of the RF during unperturbed walking is generally dissipation of mechanical energy (24) and this role can amplified or attenuated depending on the timing of the perturbation.

When interpreting our findings, two important technical limitations should be considered. The first limitation is that RF fascicle lengths measured using the custom tracking application were not validated against gold-standard measurement tools such as diffusion tensor imaging or extended field of view techniques (62,63). Further, comparison against hand-tracked ultrasound images may not serve as a high-fidelity standard because 2D ultrasound images cannot capture the complex 3D shape changes of the muscle (64), high frame rate ultrasound imaging still necessitates the extrapolation of fascicle lengths from the imaged section of the muscle, and manual tracking of a single fascicle would likely have more variability than tracking all fascicles within the imaged section of the RF. We are confident in our tracking application because 1) the selection of snippets used to calculate a representative fascicle was similar to that of a previously validated application (17), 2) only snippets within a physiological range of pennation angles were used to estimate a representative fascicle, 3) the thickness of the muscle was calculated from hand-tracked aponeuroses for each frame, 4) for within-subject designs, our extrapolation method has been shown to be acceptable for estimating normalized fascicle lengths of other quadriceps muscles (65). Further, the range of RF fascicle strains we observed during unperturbed walking (0.8 to 1.08 of fascicle slack length) are more physiologically plausible than the range reported from simulated RF fiber length during walking at 1.25 m s^-1^ (0.7 to 1.4 of optimal fiber length; (34)), when considering active strains above 25% being potentially damaging (66). The second important limitation is that we estimated fascicle and MTU forces using a Hill-type model, which is known to have shortcomings when predicting *in-vivo* forces and energetics (19,54,55,67–69). Multiple aspects of Hill-type models have been suggested to contribute to these discrepancies, a subset of which are: 1) that Hill-type models are based on phenomenological relationships characterized at maximal activations, which may explain inaccuracies when estimating forces at physiological, submaximal levels (19,54,67), 2) classical Hill-type models have independent length, velocity, and activation components and thus do not capture physiological relationships between length and activation (70–72) or velocity and activation (73), and 3) Hill-type models do not capture the history dependence of muscle force, which may play an important role in responses to perturbations (74–76). Despite these shortcomings, the computational tractability of Hill-type models has resulted in them becoming the state of the art for use in simulations of human movement (34,77,78). By using *in-vivo* estimates of fascicle length and activation to drive a Hill-type model, our approach represents a potential improvement over other fully computational approaches by avoiding assumptions about force sharing among quadriceps muscles and the influence of series elasticity from aponeuroses and the external tendon, both of which are particularly relevant for the RF (79,80).

To conclude, this study extended a previous investigation of the relationship between joint and leg level mechanical energetics during perturbed walking down to the level of the RF MTU and fascicles. This was accomplished by coupling in-vivo EMG and B-mode ultrasound imaging in conjunction with a Hill-type model of muscle force. We found that on the perturbed leg the RF MTU and fascicles actively lengthened during perturbations, but fascicle lengthening was limited due to shape change of the deep aponeurosis, 2) on the contralateral side the RF, particularly at the MTU level, better reflected energetic demands on the hip as opposed to the knee and leg levels, and 3) on the perturbed leg the RF acted to shuttle energy across the leg to fulfill the overall leg-level demand. We anticipate future work will use such multi-scale approaches to better understand how changes in muscle-level dynamics explain altered joint and leg level demands during use of assistive devices (36), and following injury (81). Further, muscle-level measurements coupled with models of physiological sensory organs can provide a window into the signals and motor programs used by the nervous system to maintain stability during locomotion (82–84).

## 1.1 Ethics

All participants (recruitment period: 8/28/2020 - 3/6/2021) provided written informed consent and all protocols were approved by the Institutional Review Board at the Georgia Institute of Technology (Protocol H20163).

## 1.2 Data Accessibility

The ultrasound processing scripts are available at https://github.com/pgolyski/RF_US_DL and biomechanical data for all participants (N=7) are available at: https://zenodo.org/doi/10.5281/zenodo.12728475

## 1.3 Authors’ Contributions

Pawel R. Golyski: Conceptualization, Data curation, Formal analysis, Methodology, Visualization, Writing – original draft, Writing – review & editing.

Gregory S. Sawicki: Conceptualization, Funding acquisition, Writing – review & editing, Project administration, Supervision

## 1.6 Acknowledgements

The authors thank Jennifer Leestma for her development of the perturbation program and insightful discussions, in addition to Patrick Kim and Nicholas Swaich for assistance with data collection.

## Supplementary Figures

**Figure S1.**
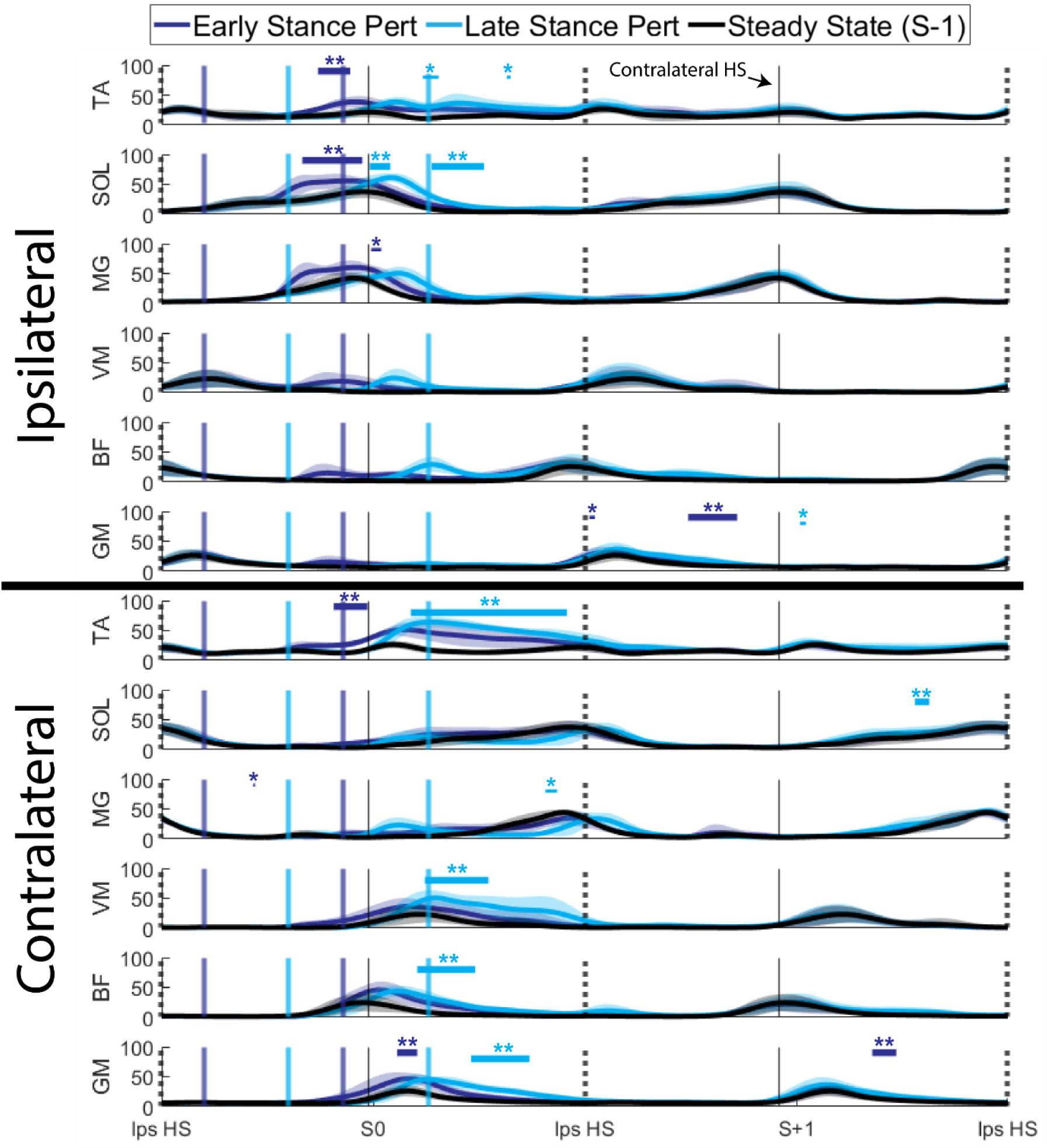
Muscle activations (% max across all trials) on both legs relative to the perturbation across all participants (n=7, 222 perturbations), time-normalized to gait cycles. TA = tibialis anterior, SOL = soleus, MG = medial gastrocnemius, VM = vastus medialis, BF =biceps femoris, GM = gluteus maximus. Shaded regions represent ±1 standard deviation. Instances when curves significantly deviated from steady state (S-1) are identified with thick horizontal lines and ** for p < 0.001, and thin horizontal lines and * for p < 0.05. Solid colored vertical lines indicate average start and end times of the perturbations. Dotted/solid black vertical lines indicate instances of ipsilateral/contralateral heelstrike, respectively. “Steady State” strides were the strides preceding the perturbed stride (S-1).

**Figure S2.**
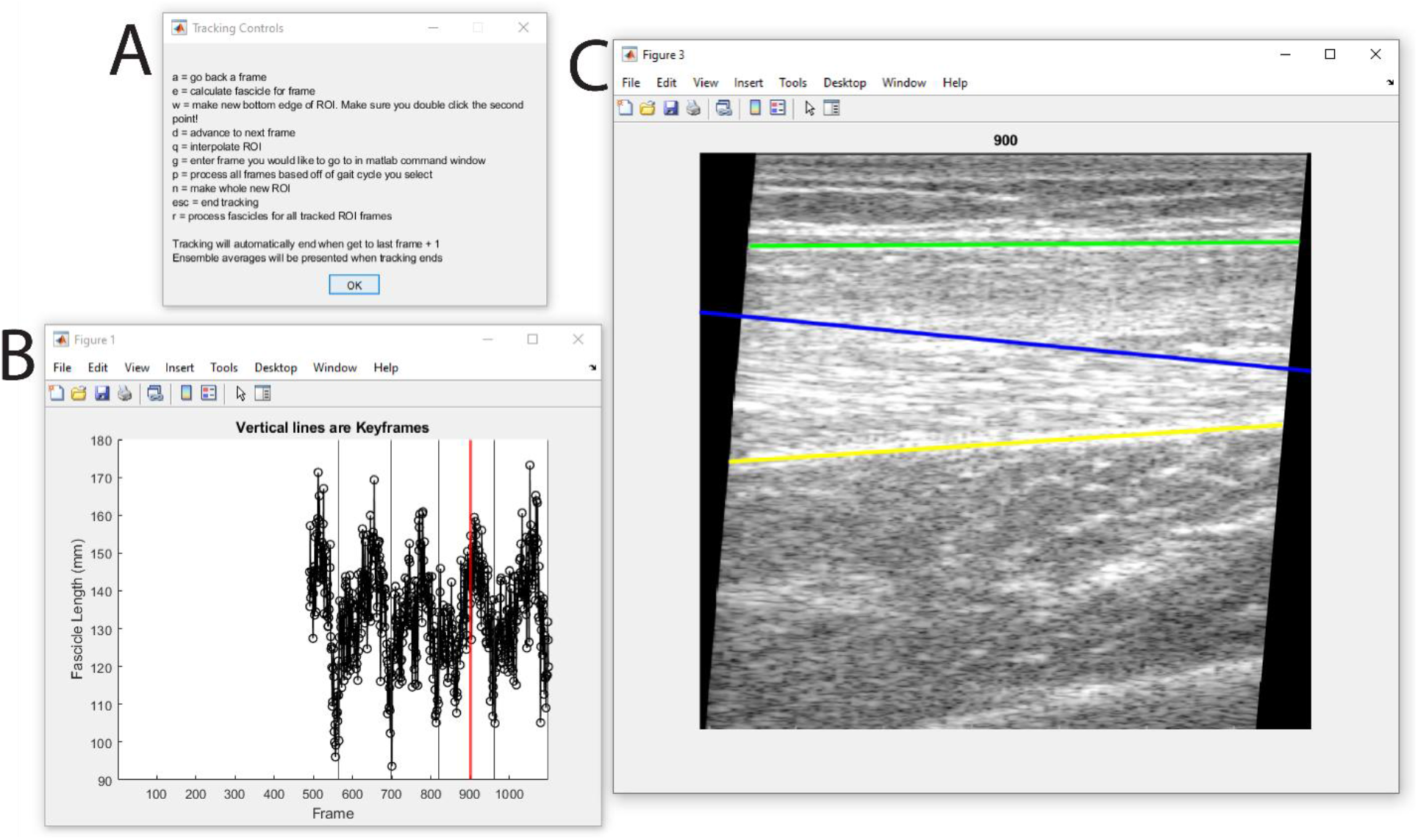
Custom fascicle tracking application user interface. A) Instructional information for operating the tracker. B) Representative fascicle length window. The vertical red line indicates the current image frame. Vertical thin black lines indicate keyframes (*e*.*g*. ipsilateral heelstrikes). C) Image window. The green line indicates the superficial aponeurosis, the yellow line indicates the deep aponeurosis, and the blue line is a graphical presentation of a representative fascicle located at the middle of the images.

**Figure S3.**
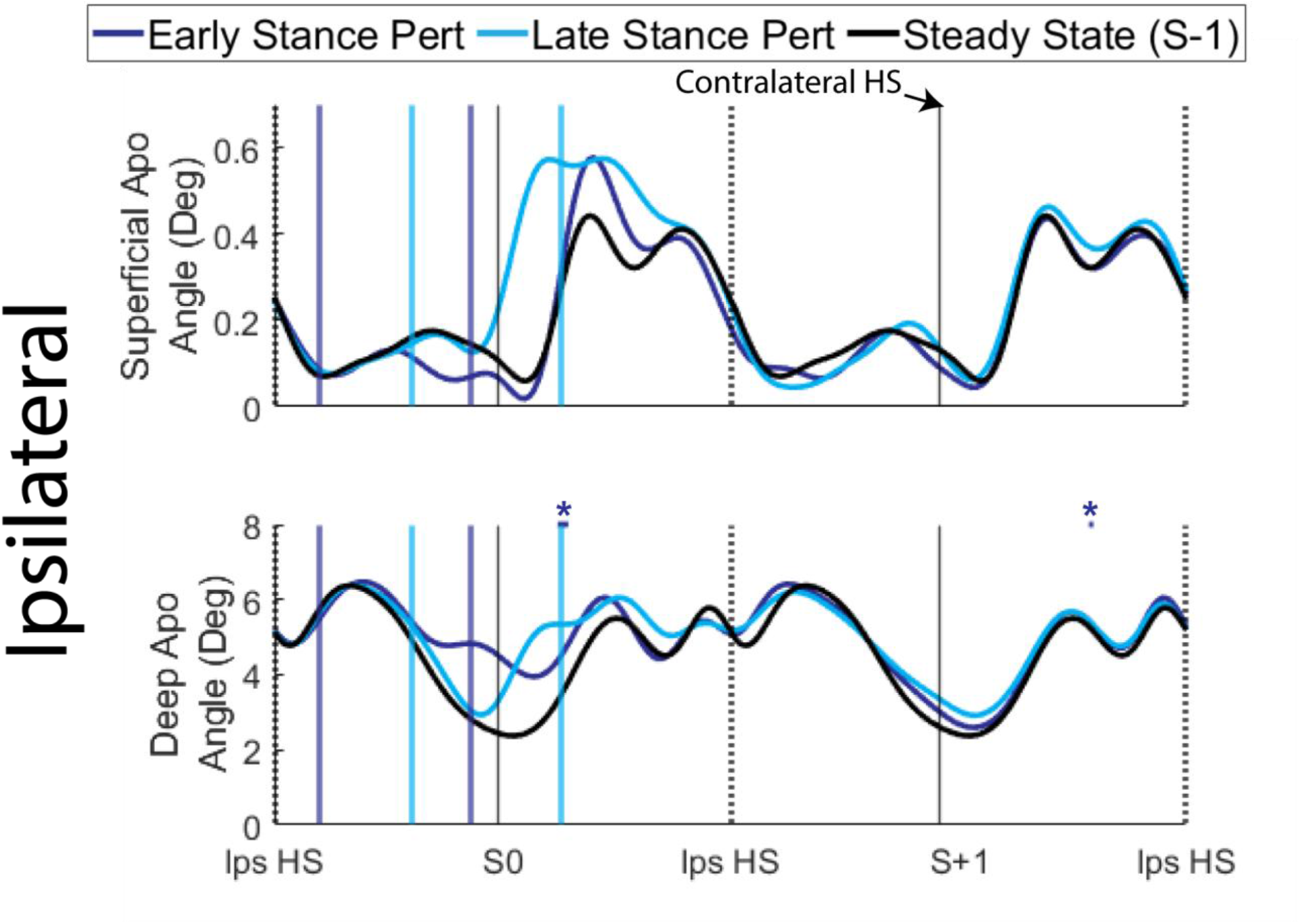
Superficial and deep aponeurosis angles relative to horizontal (positive is counterclockwise) on the ipsilateral leg relative to the perturbation across all participants (n=7, 222 perturbations), time-normalized to gait cycles. Standard deviations are omitted for clarity. Instances when curves significantly deviated from steady state (S-1) are identified with thick horizontal lines and ** for p < 0.001, and thin horizontal lines and * for p < 0.05. Solid colored vertical lines indicate average start and end times of the perturbations. Dotted/solid black vertical lines indicate instances of ipsilateral/contralateral heelstrike, respectively. “Steady State” strides were the strides preceding the perturbed stride (S-1).

